# Light color and nutrients interact to determine freshwater algal community diversity and composition

**DOI:** 10.1101/2025.02.27.640658

**Authors:** Jake A. Swanson, David A. Abdulrahman, Heather E. Bruck, Tammi L. Richardson, Jeffry L. Dudycha

## Abstract

Lakes experience a wide range of variation in resource availability; variation of essential resources such as light and nutrients can cause changes in algae community structure. Changes in light color and nutrient availability could lead to shifts in algal community diversity and composition, including shifting communities to be dominated by cyanobacteria. Due to eutrophication and brownification, lakes are experiencing increases in nutrient concentrations and shifts from blue towards red in the color of light available to algae for photosynthesis. We investigated whether differences in light color and nutrients affect algal community composition and diversity, and whether differences in light color alter the impact that nutrient levels have on algal communities. We used experimental microcosms with a fully factorial experimental design, crossing four light colors with two nutrient levels. We assessed community composition by enumerating algal taxa via light microscopy. We found that light color and the interaction between light color and nutrient availability led to large differences in community diversity, with blue light leading to the greatest diversity and broad light the lowest. Light color, nutrients, and the interaction between the two were all significant drivers of differences in community composition. Overall, we found that light color and nutrient availability interact to affect algal community diversity, composition, and cyanobacteria density. Consequences of light color are infrequently studied in aquatic ecology, but our results show that light color may need to be considered more broadly. Furthermore, our results suggest that concurrent eutrophication and brownification may yield environmental conditions favorable to cyanobacteria, including taxa that can form harmful algal blooms.

## Introduction

Lakes exhibit a wide range of natural variation of resource availability. Some of these resources, like light and nutrients, play key roles in structuring algal communities. Light as a resource can vary in two ways; in its intensity, which is how much light is available, and in its color, which is the wavelengths of light that are available. How variation of light intensity influences algal communities has been extensively studied, with changes in light intensity shown to alter community diversity and competitive outcomes among taxa (Huisman & Weissing, 1994; Litchman & Klausmeier, 2001; Flöder et al., 2002, Agawin et al., 2007). The effects of differences in light color, however, are less well known (but see Wall & Briand, 1979; Hintz et al., 2021; Stockenreiter et al., 2021; Hintz et al., 2022). Variation of the color of light available to algae for photosynthesis can increase or decrease population growth rates, depending on how closely the available colors of light match species’ absorptive pigments (et al., 2004, Luimstra et al., 2020). All photosynthetic algae share the pigment chlorophyll *a*, which absorbs specific wavelengths of blue and red light (Aguirre-Gomez et al., 2001). Niche partitioning via light color can occur because different taxa of algae have different secondary pigments such as phycobilins, carotenoids, or xanthophylls that can absorb wavelengths of light that chlorophyll *a* cannot (Stomp et al., 2008, Neun et al., 2022). Fucoxanthin, for example, is a secondary pigment unique to diatoms that has an absorption peak in the blue/green region of the light spectrum (Papagiannakis et al., 2005, Kuczynska et al., 2015). Understanding how differences in light color alter algal community composition and diversity is there important to determining how environmental variation drives community structure.

Much attention has also been given to the effects of changes in nutrient availability, particularly phosphorus and nitrogen, on algal growth (Grover 1989; Litchman et al., 2003; Sterner et al., 2004) and community diversity and composition (Guildford & Hecky, 2000, Ptacnik et al., 2008, Marzetz et al., 2020). However, differences in light color may mediate the effect of nutrient variation on algal communities. As algae form the foundation of freshwater food webs it is vital to understand whether variation in light color and nutrient availability interact to alter the diversity and composition of these communities.

In addition to natural variation of light color and nutrient availability, brownification and eutrophication are causing increases in colored dissolved organic matter (CDOM) and nutrient concentrations, respectively (Schindler et al., 2016; Hayden et al., 2019; Blanchet et al., 2022). As nutrient concentrations increase, algae that can respond quickly to these changes, such as cyanobacteria, may come to dominate algal communities (Carey et al., 2012). Increasing nutrient availability may also release some taxa from competition for nutrients, which removes a dimension through which competitive trade-offs may allow for coexistence among algal species. Additionally, taxa with smaller cell sizes are generally considered to be stronger nutrient competitors because they have a larger surface area to volume ratio which allows for more efficient nutrient uptake (Litchman & Klausmeier, 2008). Therefore, as lakes continue to be affected by eutrophication, competition for nutrients potentially may have less of a structuring effect on algal communities, which therefore may shift to become dominated by larger-celled taxa and cyanobacteria.

Brownification is causing lakes to become redder in color, with relatively less blue light but relatively more green and red light transmitted through the water column. This is due to the addition of increased amounts of terrestrially derived CDOM (Weyhenmeyer et al., 2016; Creed et al., 2018). Brownification induced shifts in the underwater light spectrum are predicted to favor the algal taxa whose pigment absorption peaks most closely match up with the colors of light available in the environment.

Differences in light color and nutrient availability have been independently shown to affect algal community composition and diversity (Watson et al., 1997, Neun et al., 2022). Exposure of marine algal communities to red light led to a decrease in species richness relative to green and broad light (Hintz et al., 2021). Blue and broad light conditions also had significantly greater amounts of green algae relative to red and green light whereas red and green light had significantly greater amounts of diatoms relative to blue and broad light (Hintz et al., 2021). In freshwater communities, brownification has been shown to lead to communities dominated by mixotrophic taxa such as cryptophytes in the genus *Cryptomonas* and the chrysophyte *Dinobryon* (Urrutia-Cordero et al., 2017). At low phosphorus levels, cryptophytes, chrysophytes, and diatoms perform well (Watson et al., 1997: Przytulska et al., 2017) whereas N-fixing taxa such as the cyanobacteria *Anabaena* and *Aphanizomenon* can tolerate low nitrogen levels (Rigosi et al., 2104). At high nutrient levels cyanobacteria are the dominant taxa (Watson et al., 1997, Heisler et al, 2008, Przytulska et al., 2017), but green algae, cyanobacteria, and dinoflagellates have also been shown to increase their biomass in response to phosphorus enrichment (Vanni & Findlay, 1990). Less attention, however, has been paid to the potential interaction between light color and nutrient availability. It is important to focus on joint variations in light color and nutrient availability because they may act synergistically to cause rapid shifts in algal community composition. Alternatively, these resource variations may act in opposition to one another, therefore leading certain taxa to be favored when light color varies but different taxa when nutrients vary. Additionally, interactions between light color and nutrient availability may create conditions that favor algae that can form harmful algal blooms (HABs). Bloom-forming taxa such as cyanobacteria are favored under high nutrient conditions (Carey et al., 2012) and have secondary pigments that can absorb green and red light (Stomp et al., 2008). Understanding how differences in light color and nutrient availability interact to alter algal communities will provide an increased understanding of how the fundamental ecology of limnetic ecosystems is predicted to change in the face of anthropogenetic driven environmental shifts. As eutrophication increases nutrient availability in lakes and brownification shifts the underwater light spectrum the likelihood of harmful algal blooms (HABs) occurring may increase. Whether variations of light color and nutrient availability create environmental conditions favorable to HAB forming algae has important implications changes in ecosystem functioning driven by eutrophication and brownification.

As such, we were interested in investigating how differences in light color and nutrient availability affect freshwater algal community diversity and composition and whether these differences can create conditions favorable for cyanobacteria. We hypothesized that environments with more wavelengths of available light (i.e., a broad spectrum) and low nutrient levels would lead to greater diversity compared to single color and high nutrient environments. In single color environments we expected the dominant taxa to be those whose photosynthetic pigments most closely match the wavelengths of available light; at high nutrient levels we anticipated cyanobacteria to dominate our communities irrespective of light due to their high fitness in nutrient rich environments. We tested our predictions with an experimental microcosm approach that used algal communities derived from lakes with different nutrient environments.

## Methods

### Field Sampling and Microcosm Set-Up

To inoculate the microcosms with phytoplankton from natural communities we sampled from two lakes in South Carolina, Lake Jocassee and Lake Murray. Lake Jocassee is an oligotrophic lake located within Devils Fork State Park, Salem, SC (34° 57′ 36″ N, 82° 55′ 10″ W), and Lake Murray is a large eutrophic reservoir spanning multiple municipalities in SC (34° 3′ 56.86″ N, 81° 19′ 44.29″ W). Sampling from both an oligotrophic and a eutrophic lake allowed us to generate a broad species pool of algae that reflected real communities yet contained taxa from both low- and high-nutrient environments. At each lake we sampled 1L of water at four different depths: surface, half Secchi depth, Secchi depth, and 1.5 times the Secchi depth. Each sample was collected with a horizontal Van Dorn sampler and was filtered through a 100 μm mesh filter to exclude zooplankton. We sampled at multiple depths to avoid biasing our species pool towards taxa that only appear at a particular depth. Both lakes were sampled on September 10^th^, 2021. Before inoculating the microcosms, we created a regional species pool of algae by combining all the samples from both lakes together for a total sample volume of 8 L.

Microcosm experimental treatments consisted of four light color treatments, blue (peak wavelength 445.5 nm), green (peak wavelength 515 nm), red (peak wavelength 656), and broad (peak wavelength 595.5 nm), crossed with two nutrient treatments, high (1 mg/L of added ammonium phosphate) and low (1 μg/L of added ammonium phosphate) for a total of eight possible experimental treatments. Microcosms were illuminated from above by LED lights.

Light intensity was set at 30 μmol/photons/m^2^s^-1^ for every light color treatment. As all microcosms were kept within the same growth chamber, we used black felt to separate light treatments from one another; this resulted in light treatments that only consisted of the color of interest (Figure S1). We replicated each treatment three times for a total of 24 microcosms. 400 mL microcosms were set up in 500 mL glass Pyrex bottles. 200 mL of the initial microcosm volume consisted of lake water drawn from the regional species pool and 200 mL consisted of a lab-made modified version Waris-Harris medium with either 1 mg/L or 1 μg/L of ammonium phosphate depending on the nutrient treatment. Microcosms were placed in a temperature-controlled growth chamber kept at 20° C with a 12h/12h day-night cycle. Microcosms were swirled to prevent cell clumping and randomly repositioned every other day.

### Microcosm Sampling

Microcosms were allowed to run for 54 days and were sampled at the start and end of the experiment (sample days: 0 and 54). 10 mL was sampled from each microcosm and preserved in ∼5% Lugol’s iodine solution for future enumeration of the algal communities. Approximately every 10 days 10 mL was sampled for whole community absorption spectra analysis. After sampling we added enough of the appropriate lab-made medium to keep the total volume of each microcosm at 400 mL. Additionally, on day 31 we replaced 200 mL in each microcosm with fresh medium.

### Algal Community Enumeration

Algal communities from day 54 samples were enumerated via light microscopy using a Zeiss AX10 inverted microscope. 1 mL of preserved sample was placed into a gridded Sedgewick-Rafter chamber with a volume of 1 mL and allowed to settle for 10 minutes. The Sedgewick-Rafter chambers were divided into a 50×20 grid, which creates 1,000 individual squares. Using data from a pilot experiment, we determined the ratio of algal cells found on the perimeter versus the interior of a Sedgewick-Rafter chamber and used that to inform our counting protocol for this experiment. For each 1 mL sample, we counted in sets of seven squares where six of the squares were interior and one was perimeter. We counted until we hit 100 algal cells in a sample or counted 28 Sedgewick-Rafter squares, whichever came first. We chose 28 as the maximum total number of squares to count because it should allow us to detect 90% or more of all species present (McAlice, 1971). If 100 algal cells were noted in a set of seven Sedgewick-Rafter squares before the seventh square, the full set of seven squares were still all counted. Each sample at a given time point was counted in triplicate, yielding nine counts per treatment (3 count replicates x 3 microcosm replicates). Count data was converted to cell density (cells/mL) by dividing 1,000 by the number of squares counted, multiplied by the number of cells counted of a particular taxon. Algae were identified to the finest taxonomic level possible with light microscopy.

### Statistical Analysis

All analyses were done in R version 4.2.2 (R Core Team). Algal community diversity at the genus level was assessed by comparing three diversity indices (Shannon, Simpson, and Inverse Simpson) across all treatments. The effects of light color, nutrient level, and the interaction between the two, were analyzed on each diversity index with a linear mixed effects model that included light color, nutrient level, and their interaction as fixed effects with microcosm replicate and count replicate nested within microcosm replicate, as random effects. We then used an ANOVA on our model output to test for the effects of light color, nutrient level, their interaction on diversity. Differences between Shannon diversity among treatments were visualized with boxplots. Here, we report and show results only for the Shannon index as analyses for the other indices were qualitatively equivalent, with the exception of nutrients having on significant effect on Simpson diversity, and are therefore reported in the supplemental information (Tables S1, S2, S3, S4). Differences in community diversity were also visualized using rank-abundance curves. Rank-abundance curves were calculated using the *rankabuncomp* function in the *BiodiversityR* package (Kindt & Coe, 2005). Plots of the proportions of the top ten most abundant taxa relative to the total abundance in each treatment, with 95% confidence intervals, were generated using the *ggplot2* (Wickham, 2016) and *ggtext* (Wilke & Wiernik, 2022) packages. These plots also include the total algal abundance for each treatment as a figure label. Differences in algal community composition at the beginning (day 0) and end (day 54) of the experiment were analyzed at the taxonomic levels of class and genus via PERMANOVA in the package *vegan* (Oksanen et al., 2022). A Bray-Curtis dissimilarity matrix was generated using the count data for each treatment; PERMANOVA was run on these matrices using the function *adonis2* in the package *vegan*. Non-metric multidimensional scaling (NMDS) was done on both the class and genus level dissimilarity matrices using the *metaMDS* function in the *vegan* package. To visualize differences in community the ordinations from the NMDS for each nutrient treatment along with the convex hulls for each light color were plotted using *ggplot2* (Wickham 2016). The results at the level of class and genus were qualitatively equivalent so here we report and show the results at the level of genus; results for the class level can be found in the supplemental information.

We tested for differences in algal density at the class level (cryptophytes, cyanobacteria, diatoms, dinoflagellates, green algae, and class unknown). by using a linear mixed-effects model and ANVOA on the linear model output. Our model had light color, nutrient level, and their interaction as fixed effects with count replicate nested within microcosm replicate as a random effect. We also looked at differences between filamentous and non-filamentous cyanobacteria density using the same linear mixed effect model and ANOVA process, with filamentous cyanobacteria density and non-filamentous cyanobacteria density each having their own model. We visualized differences in algal class density and cyanobacteria type density using boxplots made with *ggplot2*.

## Results

### Algal Community Diversity

The blue low nutrients treatment was the most diverse whereas broad high nutrients had the lowest diversity (Figure 1). Light color and the interaction between light color and nutrient level had significant effects on algal community diversity (Table 1). In blue light, green light, and broad light low nutrient treatments were more diverse than high nutrient treatments (Figure 1). However, this pattern was strikingly reversed in red light (Figure 1). High nutrient microcosms had a greater density of algae in all light treatments except for red light (Appendix S1: Figure S2). Including microcosm replicate and count replicate as random effects did not improve model performance for any diversity index (Table 2, Tables S1, S3).

**Figure 1:**
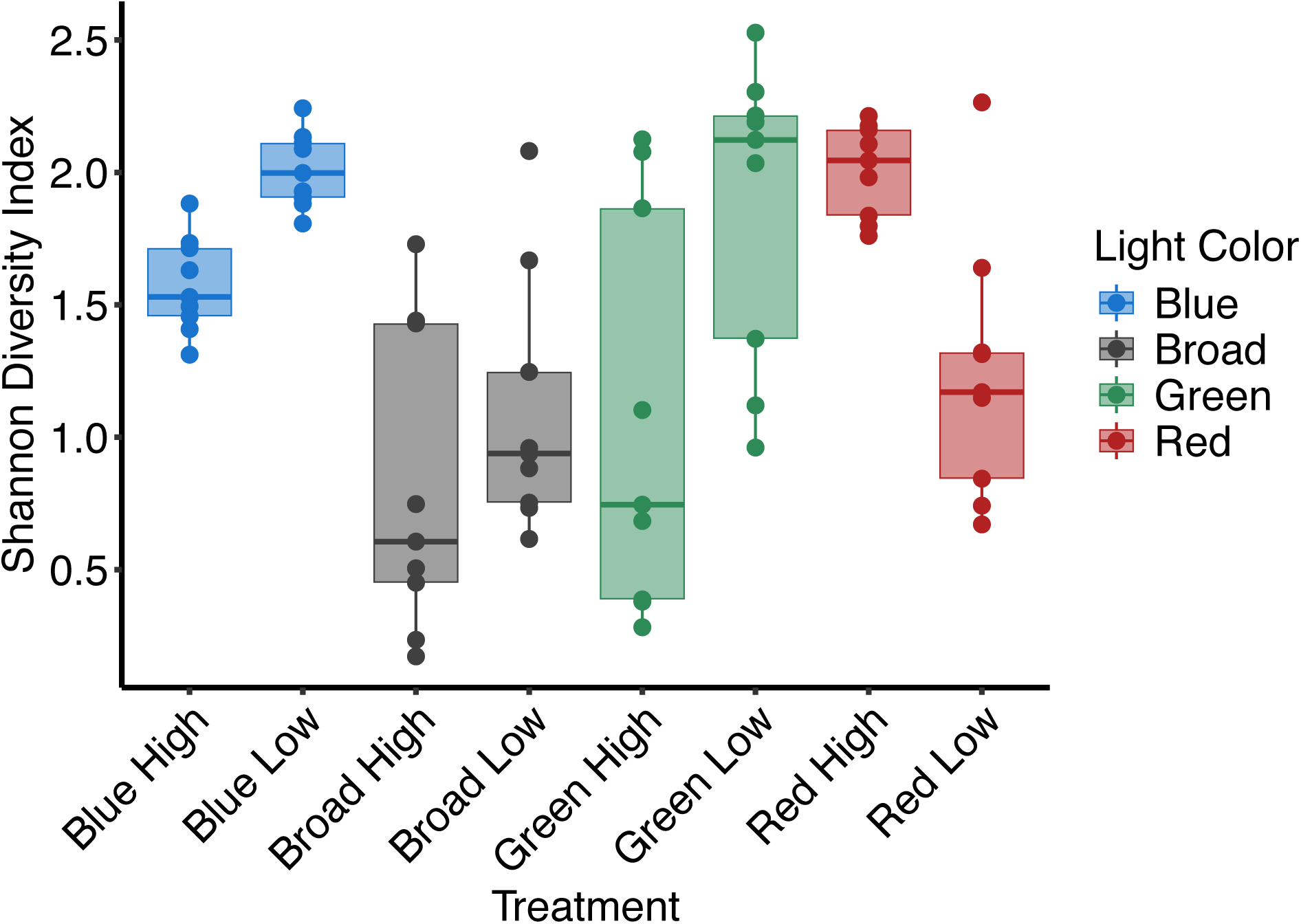
Shannon diversity index values at the level of genus for each count replicate of all microcosms for each treatment. Point and box color corresponds to each light color treatment, with black representing the broad light treatment.

**Table 1:** Output from the Shannon diversity linear mixed effects model.

| Term | Estimate | Std. Error | Df | *t* | *P* |
| --- | --- | --- | --- | --- | --- |
| Intercept | 1.57 | 0.16 | 42.78 | 9.86 | **1.39 x 10<sup>-12</sup>** |
| Broad | -0.76 | 0.22 | 62 | -3.43 | **0.0011** |
| Green | -0.50 | 0.22 | 62 | -2.26 | **0.027** |
| Red | 0.43 | 0.22 | 62 | 1.96 | 0.054 |
| Low Nutrient | 0.44 | 0.22 | 62 | 1.97 | 0.053 |
| Broad x Low Nutrient | -0.15 | 0.31 | 62 | -0.49 | 0.63 |
| Green x Low Nutrient | 0.36 | 0.31 | 62 | 1.16 | 0.25 |
| Red x Low Nutrient | -1.21 | 0.31 | 62 | -3.86 | **0.00027** |

**Table 2:**
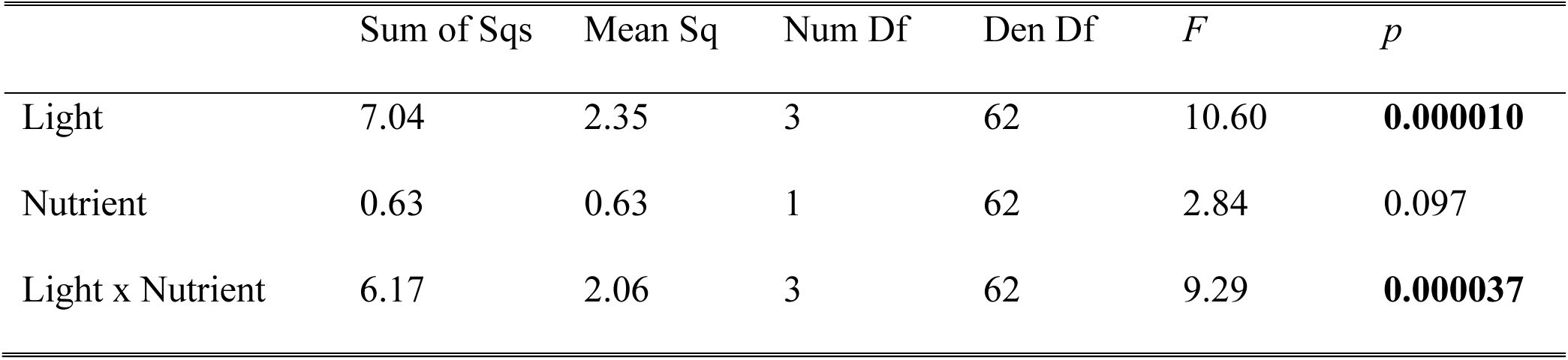
ANOVA summary statistics for the effects of light color, nutrient level, and their interaction on algal diversity calculated via the Shannon diversity index.

|  | Sum of Sqs | Mean Sq | Num Df | Den Df | *F* | *p* |
| --- | --- | --- | --- | --- | --- | --- |
| Light | 7.04 | 2.35 | 3 | 62 | 10.60 | **0.000010** |
| Nutrient | 0.63 | 0.63 | 1 | 62 | 2.84 | 0.097 |
| Light x Nutrient | 6.17 | 2.06 | 3 | 62 | 9.29 | **0.000037** |

### Algal Community Composition

Algal communities across treatments were not significantly different at the level of genus on day 0 (Figure 2, Table 1). Modest, but statistically significant differences were detectable at the level of class (Table SX); these differences did not carry-over to the end of the experiment. At day 54, at the genus level we found that light color, nutrients, and the interaction between the two led to significantly different algal communities across treatments. (Figure 2, Table 2). Light color, nutrient level, and the interaction between the two all had significant effects on community composition at the end of the experiment (Table 2). We saw that differentiation by light color was more obvious in our low nutrient microcosms, with those communities showing greater dissimilarity than the high nutrient microcosms (Figure 2). For six out of the eight treatments (broad/high, broad/low, red/high, red/low, green/high, green/low) the most dominant taxon was a cyanobacterium, whereas both blue light treatments were dominated by a green alga (Figure S2). Broad/high and red/high were dominated by the filamentous cyanobacteria (*Jaaginema* sp. and *Pseudanabaena* sp., respectively) while broad/low, red/low, green/high, and green/low were all dominated by coccoid cyanobacteria in the genus *Aphanocapsa* (Appendix S1: Figure S2). In contrast, the most abundant taxa in the blue/high and blue/low treatments were green algae (Appendix S1: Figure S2).

**Figure 2:**
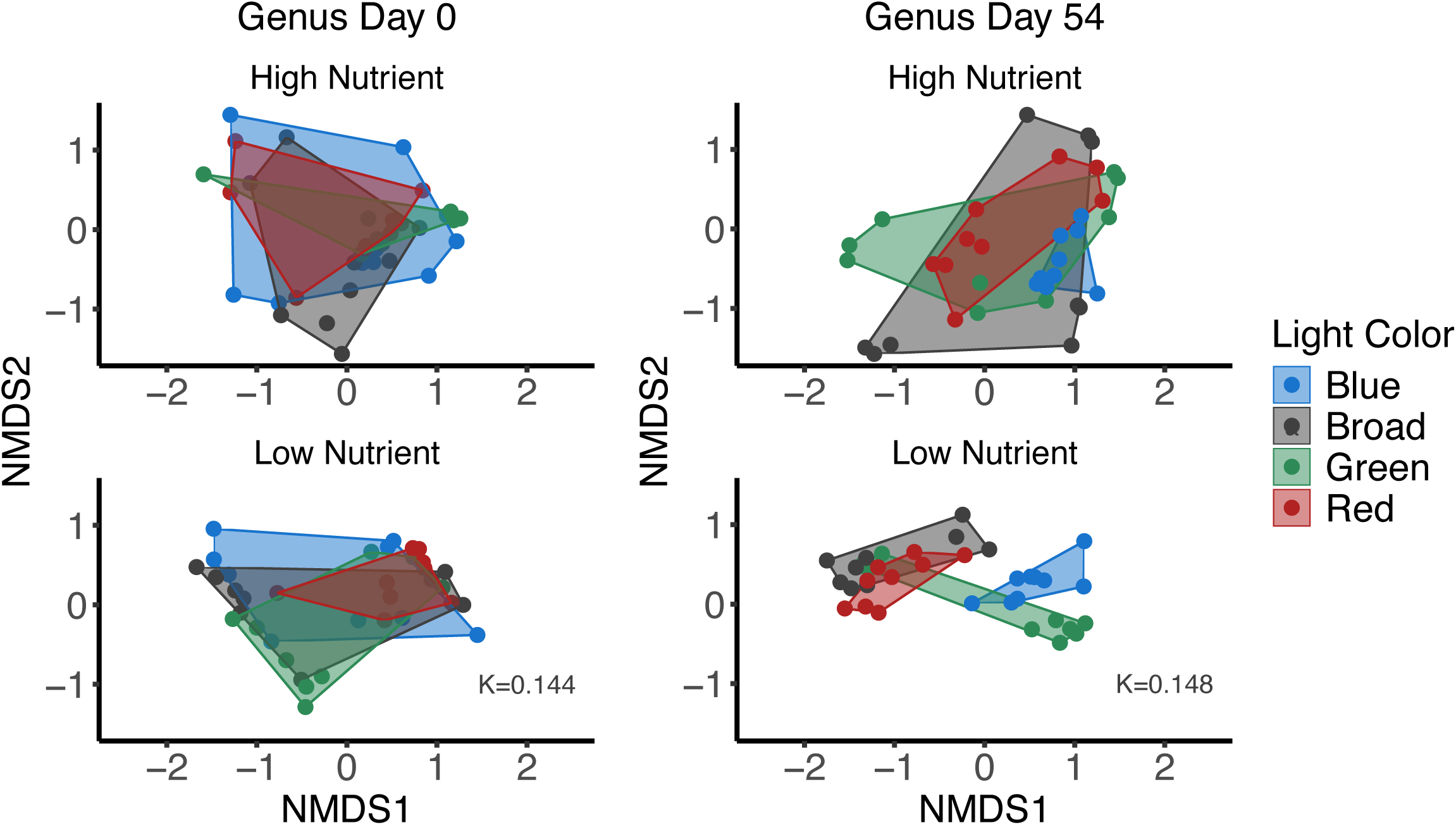
NMDS ordination plots for algal communities at the level of genus at the start (day 0) and end (day 54) of the experiment from each light color treatment at each nutrient level. Points represent the community from a count replicate for each microcosm replicate.

### *Algal Class* Density

Algal density responses to light color, nutrient level, and their interaction were class specific (Figure 2, Table 3 & Tables S-). There were no significant effects on cryptophyte density, while light color alone was a significant driver of differences in density for cyanobacteria and algae of unknown class, nutrient level alone was significant for green algae, and light color, nutrient level, and their interaction was significant for diatoms and dinoflagellates (Table SX). Cyanobacteria density was lowest in blue light across nutrient levels, and relatively high for all other colors at low nutrient levels and broad and red light at high nutrient levels (Figure 3). Green algae had greater density low nutrient conditions than high nutrient conditions regardless of light color (Figure 3). Regardless of nutrient level diatom density was lowest in broad light. At low nutrient levels it was highest in blue light and at high nutrient levels it was highest in green light (Figure 3). Green algae density trends were dependent on nutrient treatment. At low levels density was similar in all light colors while at high nutrient levels density was relatively high in blue and green light and low in red and broad light (Figure 3). Dinoflagellates and cryptophytes were less common in our microcosms than other groups which makes it difficult to make conclusions about differences in density across treatments. The significance of microcosm replicate as a random effect varied by algal class (Tables S).

**Figure 3:**
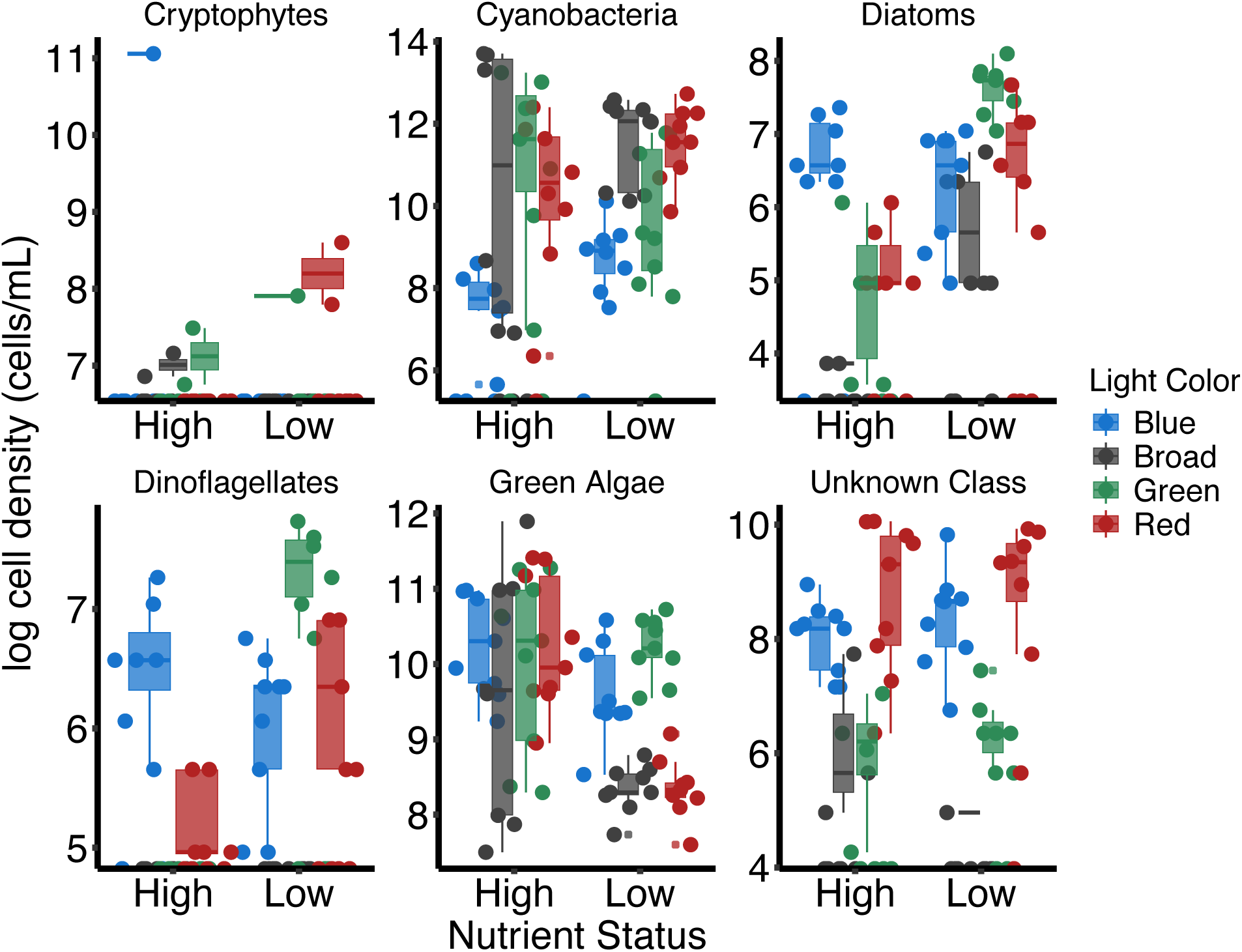
Log transformed density of major algal groups (taxonomic level of class) for each combination of nutrient and light color treatment at the end of the experiment. Point and box color corresponds to each light color treatment, with black representing the broad light treatment.

**Table 3:** Summary of PERMANOVA statistics for differences in algal community composition among treatments at the taxonomic level of genus on day 54.

| Effect | Df | Sum Of Squares | $R^2$ | $F$ | $p$ |
| --- | --- | --- | --- | --- | --- |
| Light | 3 | 4.48 | 0.19 | 6.24 | **0.001** |
| Nutrient | 1 | 2.05 | 0.08 | 8.57 | **0.001** |
| Light x Nutrient | 3 | 2.26 | 0.09 | 3.16 | **0.001** |
| Residual | 64 | 15.30 | 0.63 |  |  |
| Total | 71 | 24.09 | 1 |  |  |

**Table 4:** ANOVA summary statistics for the effects of light color, nutrient level, and their interaction on cyanobacteria density from the cyanobacteria density linear mixed-effects model.

|  | Sum of Sqs | Mean Sq | Num Df | Den Df | *F* | *p* |
| --- | --- | --- | --- | --- | --- | --- |
| Light | $3.81 \times 10^{11}$ | $1.27 \times 10^{11}$ | 3 | 62 | 5.17 | **0.0030** |
| Nutrient | $2.51 \times 10^{10}$ | $2.5 \times 10^{10}$ | 1 | 62 | 1.02 | 0.32 |
| Light x Nutrient | $1.12 \times 10^{11}$ | $3.72 \times 10^{10}$ | 3 | 62 | 1.51 | 0.22 |

**Table 5:** ANOVA summary statistics for the effects of light color, nutrient level, and their interaction on filamentous cyanobacteria type from the cyanobacteria type linear mixed-effects model.

|  | Sum of Sqs | Mean Sq | Num Df | Den Df | *F* | *p* |
| --- | --- | --- | --- | --- | --- | --- |
| Light | 4.19 x 10<sup>10</sup> | 1.40 x 10<sup>10</sup> | 3 | 266.74 | 2.69 | **0.047** |
| Nutrient | 1.50 x 10<sup>10</sup> | 1.50 x 10<sup>10</sup> | 1 | 266.98 | 2.89 | 0.090 |
| Light x Nutrient | 4.42 x 10<sup>10</sup> | 1.47 x 10<sup>10</sup> | 3 | 266.74 | 2.84 | **0.039** |

**Table 6:** ANOVA summary statistics for the effects of light color, nutrient level, and their interaction on non-filamentous cyanobacteria type from the cyanobacteria type linear mixed-effects model.

|  | Sum of Sqs | Mean Sq | Num Df | Den Df | *F* | *p* |
| --- | --- | --- | --- | --- | --- | --- |
| Light | $7.71 \times 10^9$ | 2570298380 | 3 | 541.6 | 1.91 | 0.13 |
| Nutrients | $2.83 \times 10^8$ | 282909679 | 1 | 543.97 | 0.21 | 0.65 |
| Light x Nutrients | $1.13 \times 10^{10}$ | 3762684761 | 3 | 541.6 | 2.80 | **0.039** |

### *Cyanobacteria Type* Density

Light color and the light color and nutrient interaction had significant effects on the density of filamentous cyanobacteria across treatments while nutrient level alone did not have an effect. For color and nutrient interaction. Filamentous cyanobacteria density was highest in the broad light high nutrient treatment and lowest in the blue light high nutrient treatment (Figure 4). At low nutrient levels red light had the highest filamentous cyanobacteria density and blue and green light had the lowest (Figure 4). For non-filamentous cyanobacteria red light low nutrients had the highest density and blue light high nutrient had the lowest density (Figure 4). At high nutrient levels green light had the highest non-filamentous density while at low nutrient levels blue and green light had the lowest levels of density (Figure 4).

**Figure 4:**
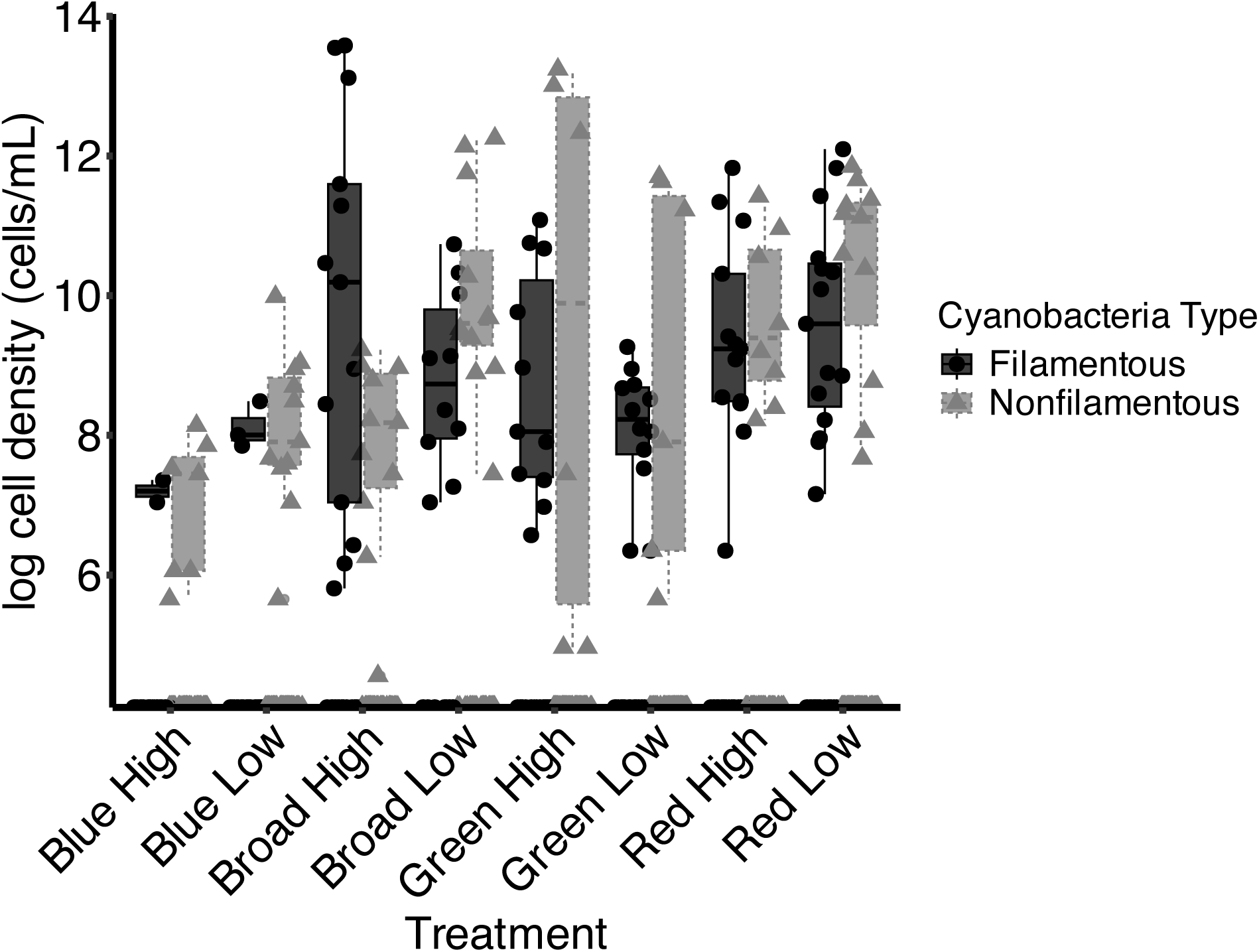
Log transformed density of filamentous and non-filamentous cyanobacteria for each treatment at the end of the experiment. Point shape and box color refers to filamentous (dots, black) or non-filamentous cyanobacteria (triangles, grey).

## Discussion

### Effects of light color and nutrients on algal diversity

Light color alone, and the interaction between light color and nutrients, had large effects on algal diversity. We expected stronger niche partitioning, and therefore higher diversity, in low nutrient conditions because algae would be competing for a greater number of scarce resources, which historically has been shown to led to increased niche partitioning (Hutchinson 1961, Tilman 1980, Burson et al., 2018). Interestingly, nutrient level alone did not influence diversity, which was unexpected given that previous work demonstrates that nutrient levels do impact freshwater algal diversity (Delgado-Molina et al., 2008; Burson et al., 2019).We expected broad light to lead to the most diverse communities because more wavelengths of light would be available compared to the single-color environments, and prior work has shown that species can coexist in broad light environments (Stomp et al., 2004, Stomp et al., 2008, Burson et al., 2019). In our microcosms, however, broad light led to the lowest levels of diversity, whereas blue light led to the highest levels of diversity while the effects of green and red light on diversity was dependent on the nutrient level of the microcosm. We predicted that, in the broad light treatment, multiple colors of light being available to the algal community would allow for taxa with unique pigment composition, such as cryptophytes, cyanobacteria, and diatoms, to coexist via niche partitioning of the light spectrum (Burson et al. 2019; Hintz et al., 2021). Instead, our results instead show that both broad high nutrient and broad low nutrient treatments were dominated by cyanobacteria, specifically the filamentous *Jaaginema* and the non-filamentous *Aphanocapsa*, respectively (Figure 3, Appendix S1: Figure S2). These cyanobacteria may have outcompeted other algae, which led to low community diversity. The high amounts of diversity seen in the blue light treatments is contrary to our hypothesis regarding light color niche differentiation. This, however, may be explained by the fact that all algal groups have chlorophyll *a*, which absorbs blue light. Multiple algal taxa, such as green algae and diatoms, can use blue light efficiently (Gorai et al., 2014, Luimstra et al., 2019, Hintz et al., 2021) whereas cyanobacteria experience decreases photosynthetic efficiency in blue light (Luimstra et al., 2018, Luimstra et al., 2019) which may have prevented them from dominating the blue light treatments as they did in other light colors.

### Effects of light color and nutrients on algal communities

Light color has been shown to impact algal community composition, with blue light environments favoring green algae (Gorai et al., 2014; Luimstra et al., 2020). This corresponds with our results that show green algae having relatively high density in blue light regardless of nutrient levels, which led to the blue light treatments being the only ones where the dominant species was not a cyanobacterium (Figure 3). Cyanobacteria, meanwhile, have been shown to be inefficient at exploiting blue light due to the underlying physiology of their photosystems (Luimstra et al., 2018), which may explain why cyanobacteria were not dominant in our blue light treatments. In green and red light, which taxa benefited was dependent on whether their pigment absorption peaks matched the colors of light available in the environment. Taxa with phycobilins, like some cyanobacteria and cryptophytes, have the secondary pigments needed to perform well in green and red-light environments (Stomp et al., 2004; Stomp et al., 2008; Heidenreich & Richardson, 2020). Cyanobacteria had high levels of density in green and red light at high nutrient levels and in red light at low nutrient levels (Figure 3), which tracks with our understanding of cyanobacteria photophysiology.

Our cryptophyte data showed that when present, cryptophyte density was highest in blue light high nutrient conditions which is contrary to our expectations given their pigment composition. Cryptophytes have secondary pigments that can absorb green or red light; therefore, we expected them to be present in greater densities in red or green light treatments.

The blue light, high nutrient treatment did have the least cyanobacteria by density, so cryptophytes may have benefited via release from competition with cyanobacteria.

We also did not observe high levels of diatom abundance in our microcosms in any treatments, most likely due to their unique need for silica to form their cell walls. Our lab-made algal medium did not include silica because we were concerned about artificially inflating diatom growth rates relative to other taxa through the addition of silica in the microcosms. The lack of silica likely prevented diatoms from reaching greater abundances in our experimental communities.

We saw the highest levels of observed dinoflagellate density in our green light low nutrient treatment and the lowest levels in red light high nutrients (Figure 3). We believe it is difficult to draw broad conclusions on the effects of our treatments on dinoflagellates for two reasons. First, we did not find many dinoflagellates in our communities, which led to relatively less data compared to other taxa. Second, dinoflagellates are not strict autotrophs and can often be mixotrophic (Taylor et al., 2008). Obtaining energy and nutrients through consumption rather than light absorption or extracellular uptake means dinoflagellates could circumvent competitive interactions by changing their trophic status.

Our results support the idea that an increase in nutrients, particularly phosphorus, can lead to communities dominated by cyanobacteria, and potentially can cause subsequent algal blooms. The addition of nutrients to the microcosms tended to create denser communities dominated by cyanobacteria. High nutrient treatments led to more dense communities in three out of the four light colors (all except red, Figure S2). Broad, green, and red light all had a species of cyanobacteria as their most abundant species under high nutrient conditions (*Jaaginema* sp., *Aphanocapsa* sp., and *Pseudanabaena* sp., respectively) This is as we expected, since increasing nutrient availability, particularly phosphorous, has been shown to shift community composition towards cyanobacteria (O’Neil et al., 2012; Bormans et al., 2016; Jankowiak et al., 2019). Three of the four high nutrient treatments (all except blue high nutrients) had a cyanobacterium as the most dominant species (Figure S2) which is not surprising as cyanobacteria are well adapted to take advantage of nutrient increases (O’Neil et al., 2012). They can rapidly increase growth rates in high nutrient conditions, leading to the competitive exclusion of other algal species. Additionally, high nutrient levels may have reduced nutrient competition in the community which could increase competition for other resources such as light. High nutrient levels also increased total community density, which may have also reduced the amount of overall light and exacerbated competition for light.

Green algae and cyanobacteria were the most prevalent and abundant taxa in all communities (Figure SX), which confounded out initial predictions of niche partitioning by light color. We saw little evidence of niche partitioning as the top ten most abundant taxa in every treatment were either green algae or cyanobacteria, apart from two cryptophyte species in blue high nutrient communities and one cryptophyte species in red low nutrient communities (Figure SX). There is evidence that cyanobacteria are more efficient at using light at low light levels relative to other taxa (Schwaderer et al., 2011); as density in the microcosms increased, competition for light would increase, which could favor low-light adapted taxa like cyanobacteria. Additionally, cyanobacteria tend to dominate algal communities as nutrient levels, particularly phosphorous, increase (Paerl et al., 2001; Rigosi et al., 2014; Przytulska et al., 2017). Therefore, in the broad high nutrient microcosms, it is not surprising that the filamentous cyanobacterium *Jaaginema* exploited the low light high nutrient environment to dominate the algal community. In the broad light low nutrient treatment, *Aphanocapsa* may have outcompeted other taxa due to its small cell size. Smaller cells are considered stronger nutrient competitors because of the way the surface-volume ratio scales with cell size (Litchman & Klausmeier, 2008) Therefore, *Aphanocapsa* may be considered a strong nutrient competitor due to its size and may have benefited from low light conditions in the same manner as *Jaaginema*. Taken together, these traits may help explain the low levels of algal diversity seen in our microcosms, particularly in our broad light treatments.

We were also interested in the split between filamentous and non-filamentous cyanobacteria in the communities as filamentous cyanobacteria can present unique ecological challenges. While both morphologies can form harmful algal blooms and produce toxins (Posch et al., 2012l; Rigosi et al., 2014; Kurmayer et al., 2016), filamentous cyanobacteria, due to their size, can also interfere with feeding by zooplankton, particularly ecologically important taxa like *Daphnia*. (DeMott et al., 2001, Sukenik et al, 2015). Reduced zooplankton grazing can then cause shifts in lake food wed structure (Sukenik et al., 2015). Two of our treatments, broad high nutrients and red high nutrients, were dominated by the filamentous cyanobacteria *Jaaginema* sp. and *Pseudanabaena* sp., respectively Species in the genus *Pseudanabaena* can form algal blooms and produce toxin (Olvera-Ramírez et al., 2010) which may have negatively impacted other algae in community. In contrast, none of our low nutrient treatments had a filamentous cyanobacterium as their most dominant species, although *Jaaginema* was the second most abundant species in broad and red light. (Figure S2). The ability of filamentous cyanobacteria to produce toxins, form blooms, and inhibit zooplankton grazing may deleteriously impact lake ecosystems; our results imply that changes in light color and nutrient levels can create favorable environmental conditions for filamentous cyanobacteria.

### Eutrophication, brownification, and the implications for freshwater algal communities

Lakes are experiencing concurrent eutrophication and brownification which is causing the nutrient and light environments within these lakes to change (Leech et al., 2018). The light environment of browner lakes is shifted towards the green and red end of the light spectrum while eutrophied lakes have higher nutrient levels. Our results potentially indicate what the algal communities of lakes may look like in the future. Brownification shifts the underwater light spectrum towards green and red light via an increase in CDOM which absorbs blue light. Both of our blue light treatments had green algae as their most dominant taxon, whereas the green high nutrients and red high nutrients treatments were dominated by cyanobacteria, both coccoid and filamentous. The algal communities in lakes that are undergoing eutrophication and brownification may shift to resemble our experimental communities more closely, both in overall community density, diversity, and composition. More specifically, our results may point towards a future where environmental conditions in lakes are more favorable for HABs. As brownification is predicted to shift the underwater light spectrum towards red and away from blue light, while eutrophication will increase nutrient availability, we may see natural algal communities that reflect our red HP treatment. The most abundant taxon in this treatment was a species of *Pseudanabeana*, a filamentous toxic cyanobacterium. *Pseudanabeana* produces microcystin, which is a toxin that has been shown to be harmful to algae (Hu et al., 2005) and grazers such as *Daphnia* and *Ceriodaphnia* (DeMott 1999; Olvera-Ramirez et al., 2010). This potential shift to less diverse and more dense communities that are dominated by cyanobacteria may have detrimental effects on the ecology and ecosystem functioning of lakes by producing harmful toxins (Paerl et al., 2001), decreasing energy transfer through food webs (Filstrup et al., 2014), altering biogeochemical cycling (Grey et al., 2000), and reducing oxygen concentrations in the water column (Paerl, 1988). Overall, our results indicate that the environmental shifts induced by eutrophication and brownification have the potential to interact to affect freshwater algal community diversity and composition, which may negatively affect ecosystem functioning.

## Acknowledgements

We thank Eric LoPresti for comments on earlier drafts of the manuscript. JAS would also like to thank John Wehr for algae identification training. This study was supported by the National Science Foundation (NSF) Dimensions of Biodiversity program under grant #1542555 to T.L. Richardson and J. L. Dudycha and a University of South Carolina SPARC grant to JAS.

## Author Contributions

The experiment was designed by JAS, JLD, and TLR. Field work was done by JAS, HEB, and DPA. Microcosm set-up, maintenance, and sampling was done by JAS, HEB, and DPA. Cell counts were done by JAS and DPA. Statistical analyses were done by JAS. The manuscript was written by JAS with all authors contributing to the editing process.

