## Supplemental Information for "Light color and nutrients interact to determine freshwater algal community diversity and composition"

Supplemental appendix to main manuscript titled: “Light color and phosphorus availability interact to determine freshwater algal community diversity and composition”

Authors: Jake A. Swanson

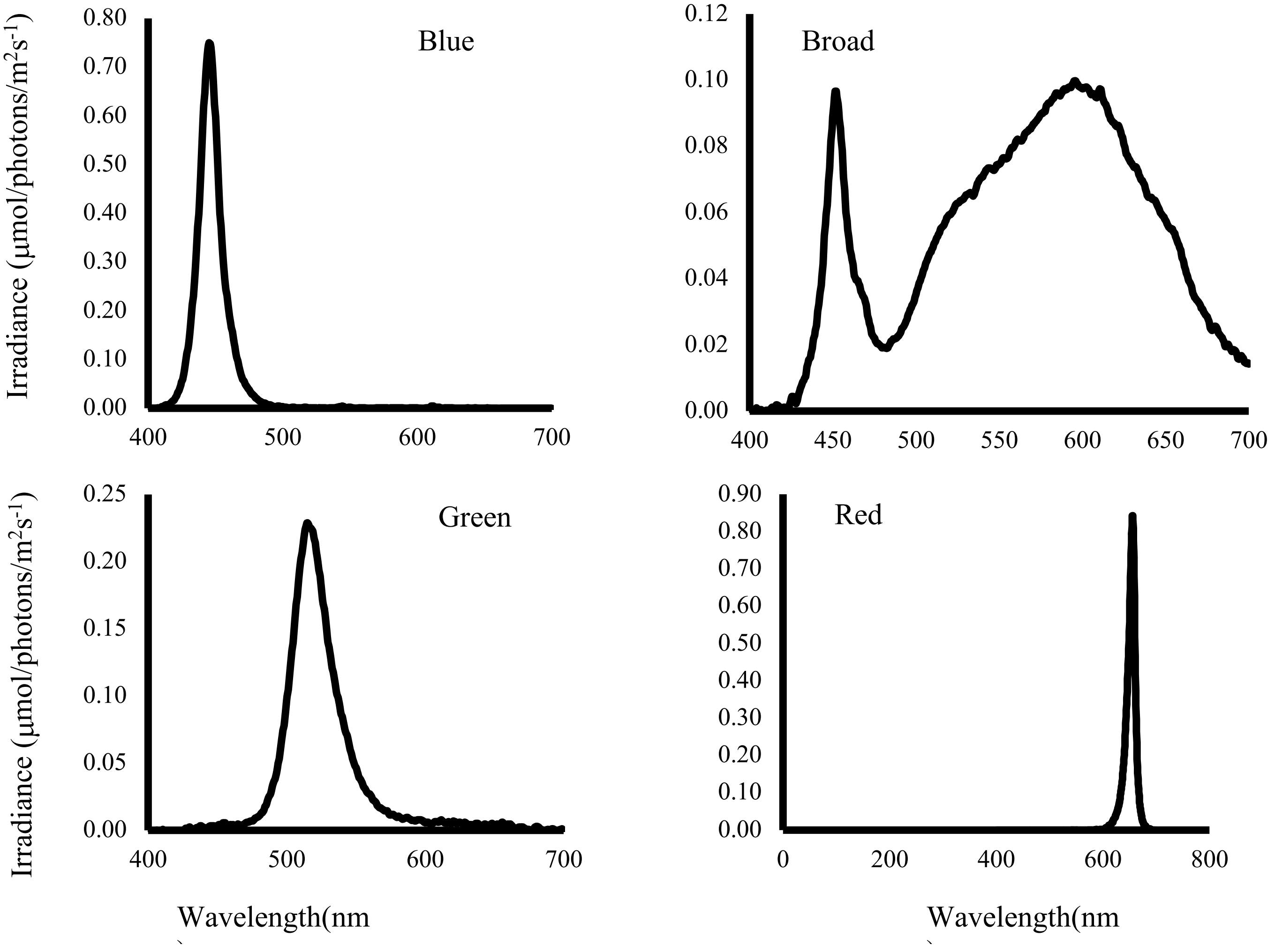


Figure S1: Light spectra for each of the four light treatments in the growth chamber.

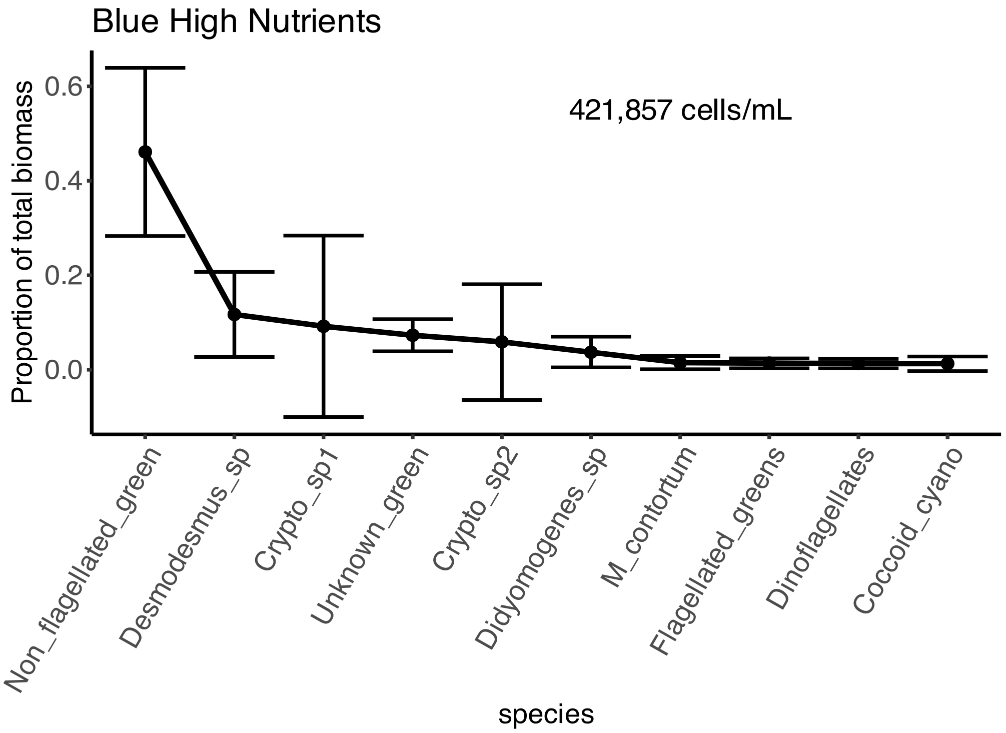


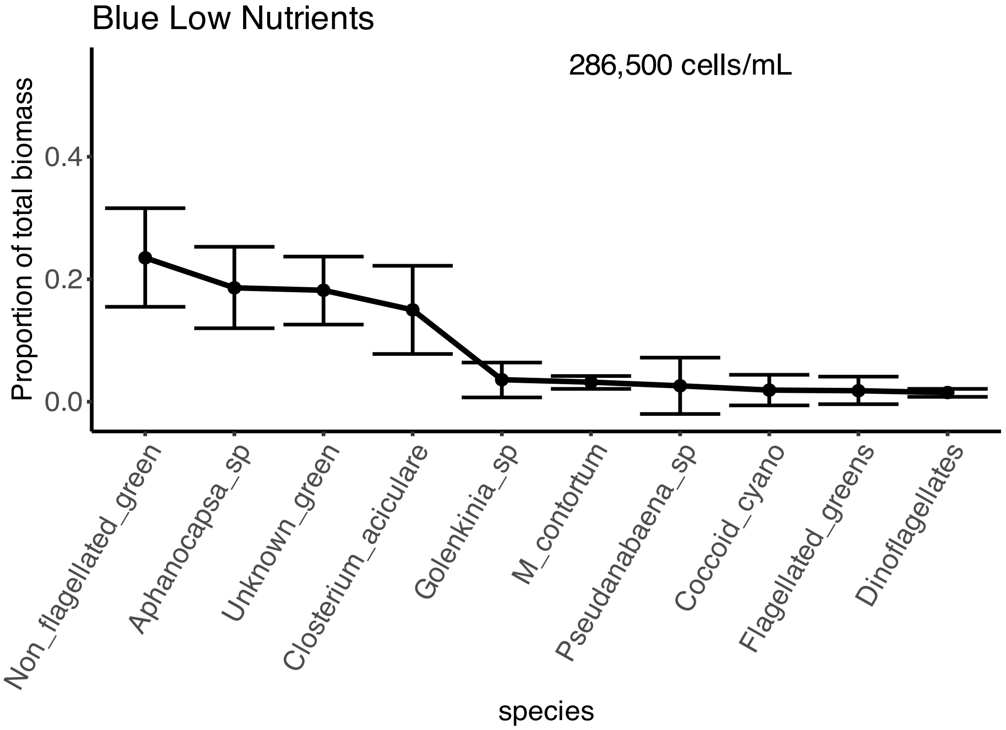


Figure S2: Rank abundance curves for the proportions of top ten most abundant algal taxa relative to total community biomass across microcosm treatments. Points and error bars represent the mean proportion and the 95% confidence interval, respectively, for each taxon across all count replicates for a microcosm.

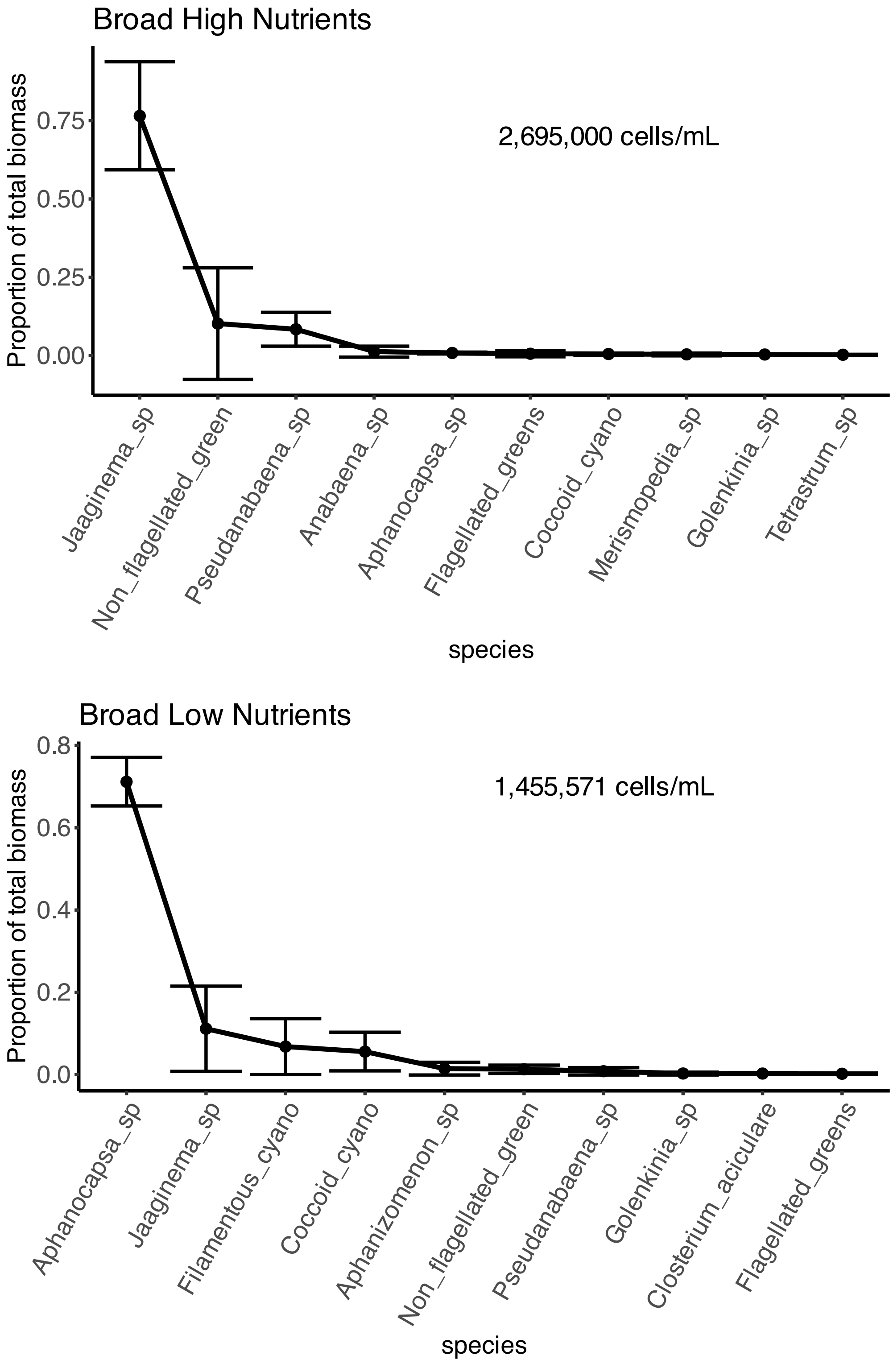


Figure S2 cont'd

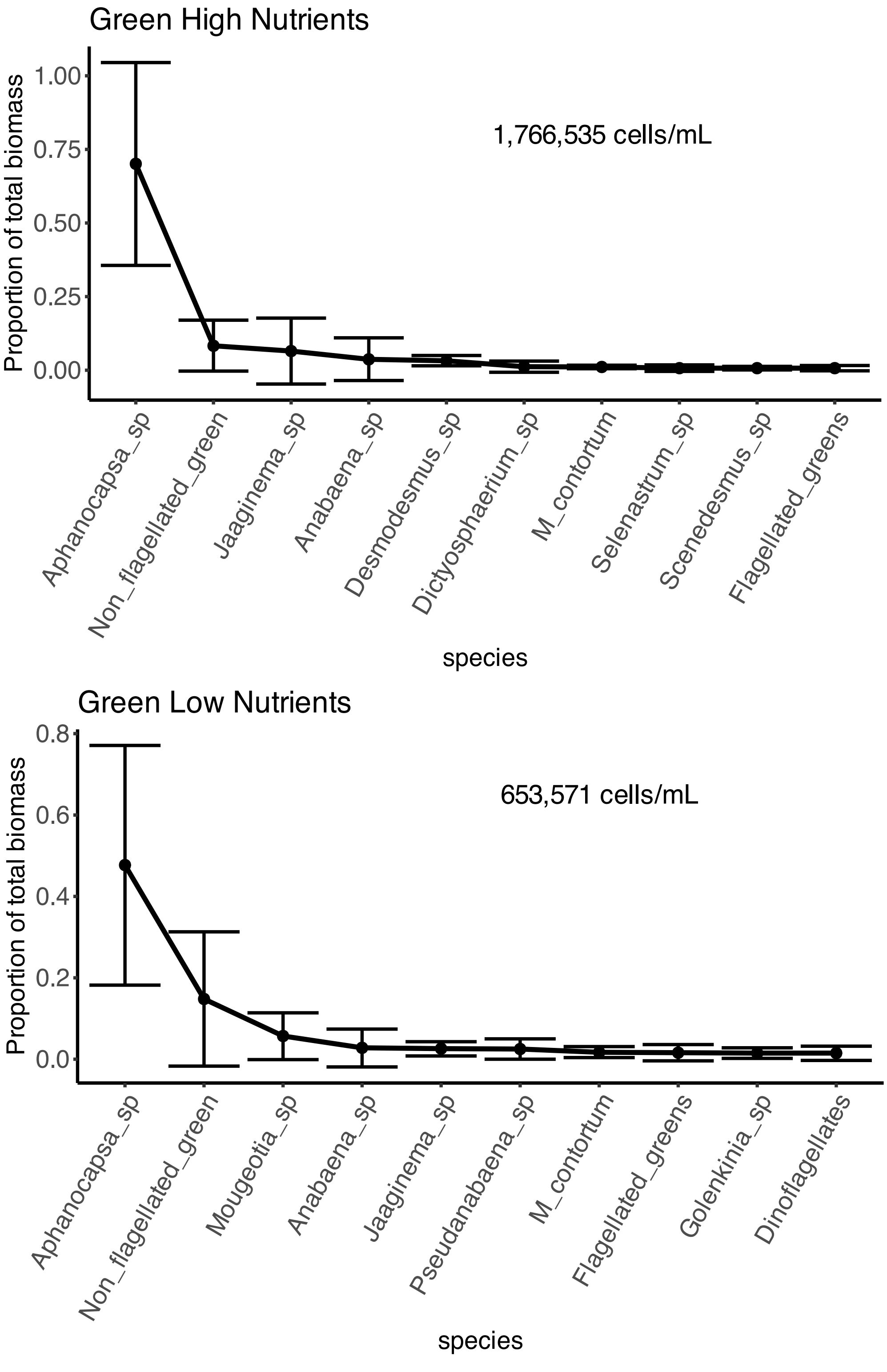


Figure S2 cont'd

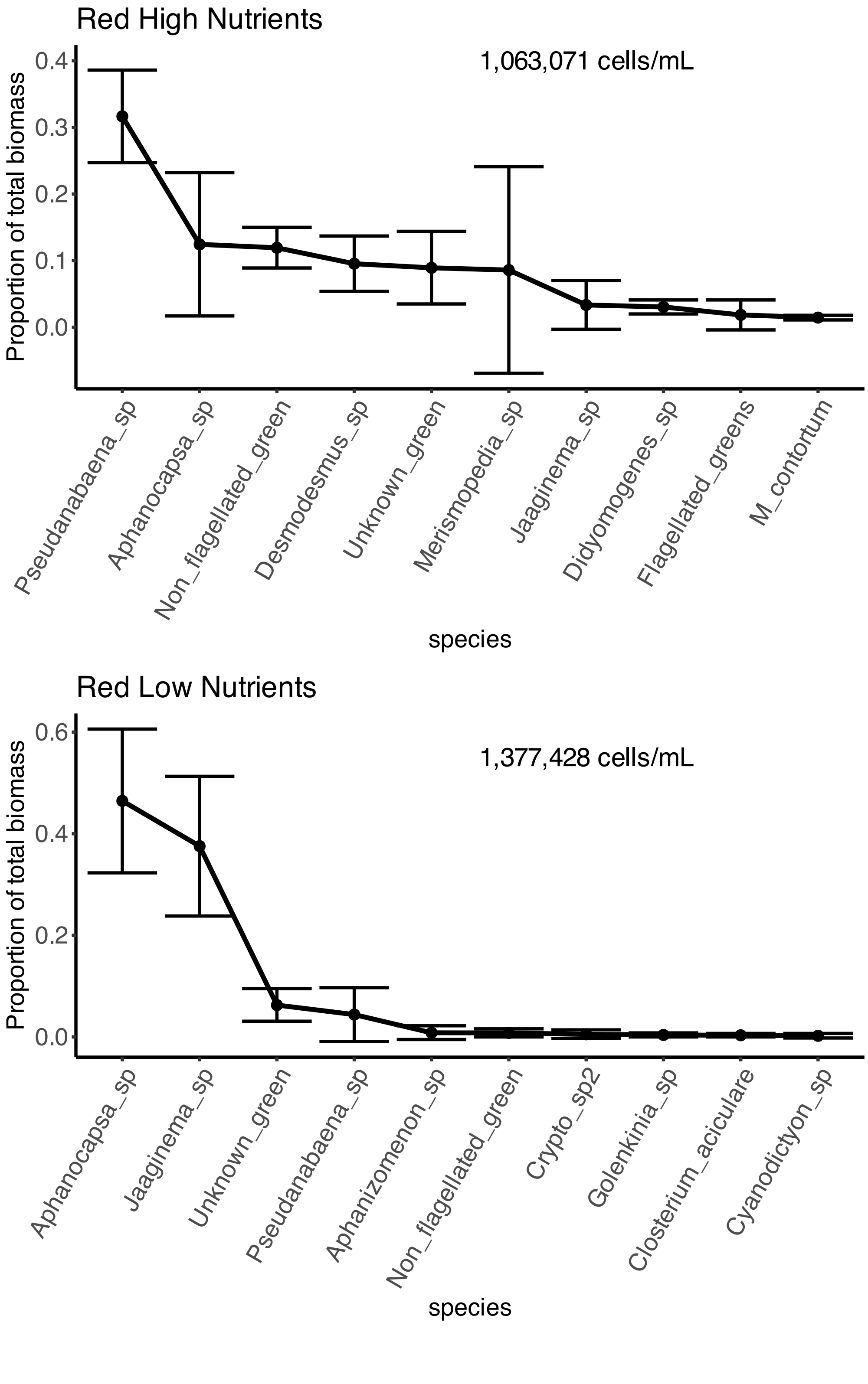


Figure S2 cont'd

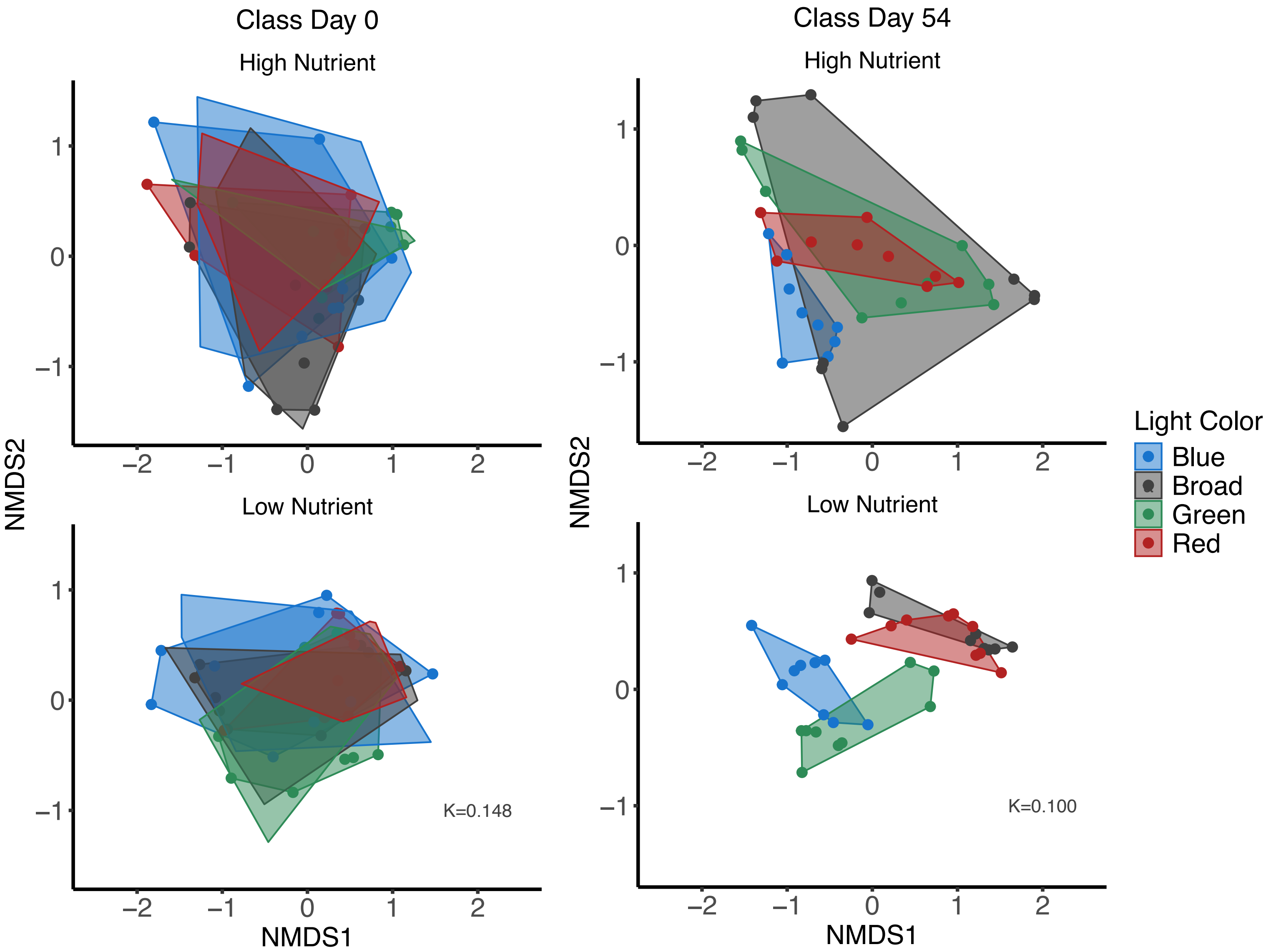


Figure S3: NMDS ordination plots for algal communities at the level of class from each treatment. Points represent the community from a count replicate of a microcosm.

Table S1: ANOVA summary statistics for the effects of light color, phosphorus level, and their interaction on algal diversity calculated via the Simpson diversity index.

| | Sum of Sqs | Mean Sq | Num Df | Den Df | *F* | *p* |
|---|---|---|---|---|---|---|
| Light | 1.012030 | 0.337343 | 3 | 62 | 11.22697 | **0.00000559** |
| Phosphorus | 0.109567 | 0.109567 | 1 | 62 | 3.64644 | **0.061** |
| Light x Phosphorus | 0.602274 | 0.200758 | 3 | 62 | 6.68134 | **0.000557** |

Table S2: ANOVA-like table for the random effects of microcosm replicate and count replicate in our Shannon diversity linear mixed effects model.

| Model | Number of parameters | log likelihood | AIC | Likelihood ratio | Df | *p* |
|---|---|---|---|---|---|---|
| No random effects | 11 | -51.6053 | 125.211 | | | |
| Random effect: Count rep nested in microcosm | 10 | -51.6053 | 123.211 | 0 | 1 | 1 |
| Random effect: Microcosm | 10 | -51.6053 | 123.271 | 0.0606426 | 1 | 0.80548 |

Table S2: ANOVA-like table for the random effects of microcosm replicate and count replicate in our Simpson diversity linear mixed effects model.

| Model | Number of parameters | log likelihood | AIC | Likelihood ratio | Df | *p* |
|---|---|---|---|---|---|---|
| No random effects | 11 | 11.7948 | -1.58957 | | | |
| Random effect: Count rep nested in microcosm | 10 | 11.7948 | -3.58957 | 0 | 1 | 1 |
| Random effect: Microcosm | 10 | 11.4327 | -2.86531 | 0.724262 | 1 | 0.39475 |

Table S3: ANOVA summary statistics for the effects of light color, phosphorus level, and their interaction on algal diversity calculated via the Inverse Simpson diversity index.

| | Sum of Sqs | Mean Sq | Num Df | Den Df | F | P |
|---|---|---|---|---|---|---|
| Light | 53.2199 | 17.73995 | 3 | 64 | 9.11641 | **0.0000415** |
| Phosphorus | 2.4137 | 2.41372 | 1 | 64 | 1.24039 | 0.269 |
| Light x Phosphorus | 74.5665 | 24.85549 | 3 | 64 | 12.77302 | **0.0000012** |

Table S4: ANOVA-like table for the random effects of microcosm replicate and count replicate in our Inverse Simpson diversity linear mixed effects model.

| Model | Number of parameters | log likelihood | AIC | Likelihood ratio | Df | *p* |
|---|---|---|---|---|---|---|
| No random effects | 11 | -120.905 | 263.81 | | | |
| Random effect: Count rep nested in microcosm | 10 | -120.905 | 261.81 | 0 | 1 | 1 |
| Random effect: Microcosm | 10 | -120.905 | 261.81 | 0 | 1 | 1 |

Table S5: Summary of PERMANOVA statistics for differences in algal community composition among treatments at the taxonomic level of genus on day 0.

| Effect | Df | Sum Of Squares | $R^2$ | *F* | *p* |
|---|---|---|---|---|---|
| Light | 3 | 1.4962 | 0.06932 | 1.7491 | **0.039** |
| Phosphorus | 1 | 0.507 | 0.02349 | 1.7782 | 0.1 |
| Light x Phosphorus | 3 | 1.3307 | 0.06166 | 1.5556 | 0.074 |
| Residual | 64 | 18.249 | 0.84553 | | |
| Total | 71 | 21.583 | 1 | | |

Table S6: Summary of PERMANOVA statistics for differences in algal community composition among treatments at the taxonomic level of class on day 0.

| Effect | Df | Sum Of Squares | $R^2$ | $F$ | $p$ |
|---|---|---|---|---|---|
| Light | 3 | 1.0585 | 0.06293 | 1.5612 | 0.116 |
| Phosphorus | 1 | 0.4147 | 0.02465 | 1.8348 | 0.109 |
| Light x Phosphorus | 3 | 0.8838 | 0.05254 | 1.3035 | 0.199 |
| Residual | 64 | 14.4643 | 0.85988 | | |
| Total | 71 | 16.8213 | 1 | | |

Table S7: Summary of PERMANOVA statistics for differences in algal community composition among treatments at the taxonomic level of class on day 54.

| Effect | Df | Sum Of Squares | $R^2$ | *F* | *p* |
|---|---|---|---|---|---|
| Light | 3 | 3.8977 | 0.21073 | 7.0831 | **0.001** |
| Phosphorus | 1 | 1.1444 | 0.06187 | 6.2391 | **0.002** |
| Light x Phosphorus | 3 | 1.7147 | 0.09271 | 3.1161 | **0.004** |
| Residual | 64 | 11.7393 | 0.63469 | | |
| Total | 71 | 18.4962 | 1 | | |

Table S8: Output from the cryptophyte biomass linear mixed effects model

| Term | Estimate | Std. Er | Dg | *t* | *p* |
|---|---|---|---|---|---|
| Intercept | 7048 | 2506 | 64 | 2.813 | 0.00652 |
| Broad | -6799 | 3544 | 64 | -1.919 | 0.05949 |
| Green | -6754 | 3544 | 64 | -1.906 | 0.06115 |
| Red | -7048 | 3544 | 64 | -1.989 | 0.051 |
| Low Nutrient | -7048 | 3544 | 64 | -1.989 | 0.051 |
| Broad x Low Nutrient | 6799 | 5011 | 64 | 1.357 | 0.17964 |
| Green x Low Nutrient | 7056 | 5011 | 64 | 1.408 | 0.164 |
| Red x Low Nutrient | 7921 | 5011 | 64 | 1.581 | 0.11891 |

Table S9: ANOVA summary statistics for the effects of light color, nutrient level, and their interaction on cryptophyte biomass.

| | Sum of Sqs | Mean Sq | Num Df | Den Df | *F* | *p* |
|---|---|---|---|---|---|---|
| Light | 142393070 | 47464357 | 3 | 64 | 0.84 | 0.477 |
| Nutrient | 46301217 | 46301217 | 1 | 64 | 0.8194 | 0.3687 |
| Light x Nutrient | 180917257 | 60305752 | 3 | 64 | 1.0672 | 0.3693 |

Table S10: ANOVA-like table for the random effects of microcosm replicate and count replicate in cryptophyte biomass linear mixed effects model

| Model | Number of parameters | log likelihood | AIC | Likelihood ratio | Df | *p* |
|---|---|---|---|---|---|---|
| No random effects | 11 | -670.8 | 1363.6 | | | |
| Random effect: Count rep nested in microcosm | 10 | -670.8 | 1361.6 | 0 | 1 | 1 |
| Random effect: Microcosm | 10 | -670.8 | 1361.6 | 0 | 1 | 1 |

Table S11: ANOVA-like table for the random effects of microcosm replicate and count replicate in our cyanobacteria biomass linear mixed effects model.

| Model | Number of parameters | log likelihood | AIC | Likelihood ratio | Df | $p$ |
|---|---|---|---|---|---|---|
| No random effects | 11 | -867.23 | 1756.5 | | | |
| Random effect: Count rep nested in microcosm | 10 | -867.23 | 1754.5 | 0 | 1 | 1 |
| Random effect: Microcosm | 10 | -870.24 | 1760.5 | 6.0306 | 1 | **0.01406** |

Table S12: Output from the cyanobacteria biomass linear mixed effects model

| Term | Estimate | Std. Er | df | t | p |
|---|---|---|---|---|---|
| Intercept | 1762 | 70670 | 7 | 0.025 | 0.980792 |
| Broad | 261700 | 73890 | 62 | 3.542 | **0.000761** |
| Green | 157500 | 73890 | 62 | 2.131 | **0.037033** |
| Red | 65520 | 73890 | 62 | 0.887 | 0.378626 |
| Low Nutrient | 5937 | 73890 | 62 | 0.08 | 0.936223 |
| Broad x Low Nutrient | -112400 | 104500 | 62 | -1.075 | 0.286337 |
| Green x Low Nutrient | -124800 | 104500 | 62 | -1.194 | 0.23702 |
| Red x Low Nutrient | 63970 | 104500 | 62 | 0.612 | 0.542671 |

Table S13: Output from the diatom biomass linear mixed effects model

| Term | Estimate | Std. Er | df | t | p |
|---|---|---|---|---|---|
| Intercept | 746.03 | 163.53 | 15.68 | 4.562 | 0.000336 |
| Broad | -735.45 | 201.36 | 62 | -3.652 | 0.000536 |
| Green | -626.98 | 201.36 | 62 | -3.114 | 0.002795 |
| Red | -603.17 | 201.36 | 62 | -2.996 | 0.003934 |
| Low Nutrient | -71.43 | 201.36 | 62 | -0.355 | 0.723993 |
| Broad x Low Nutrient | 330.69 | 284.76 | 62 | 1.161 | 0.249983 |
| Green x Low Nutrient | 2111.11 | 284.76 | 62 | 7.414 | 4.13E-10 |
| Red x Low Nutrient | 626.98 | 284.76 | 62 | 2.202 | 0.031411 |

Table S14: ANOVA summary statistics for the effects of light color, nutrient level, and their interaction on diatom biomass

| | Sum of Sqs | Mean Sq | Num Df | Den Df | *F* | *p* |
|---|---|---|---|---|---|---|
| Light | 9830215 | 3276738 | 3 | 62 | 17.959 | 1.68E-08 |
| Nutrient | 8713656 | 8713656 | 1 | 62 | 47.758 | 3.07E-09 |
| Light x Nutrient | 11722033 | 3907344 | 3 | 62 | 21.416 | 1.23E-09 |

Table S15: ANOVA-like table for the random effects of microcosm replicate and count replicate in diatom biomass linear mixed effects model

| Model | Number of parameters | log likelihood | AIC | Likelihood ratio | Df | *p* |
|---|---|---|---|---|---|---|
| No random effects | 11 | -488.52 | 999.05 | | | |
| Random effect: Count rep nested in microcosm | 10 | -488.52 | 997.05 | 0 | 1 | 1 |
| Random effect: Microcosm | 10 | -489.36 | 998.71 | 1.6606 | 1 | 0.1975 |

Table S16: Output from the dinoflagellate biomass linear mixed effects model

| Term | Estimate | Std. Er | df | t | p |
|---|---|---|---|---|---|
| Intercept | 603.17 | 145.8 | 19.21 | 4.137 | 0.000549 |
| Broad | -603.17 | 184.83 | 62 | -3.263 | 0.001793 |
| Green | -603.17 | 184.83 | 62 | -3.263 | 0.001793 |
| Red | -492.06 | 184.83 | 62 | -2.662 | 0.009874 |
| Low Nutrient | -126.98 | 184.83 | 62 | -0.687 | 0.494623 |
| Broad x Low Nutrient | 126.98 | 261.39 | 62 | 0.486 | 0.628817 |
| Green x Low Nutrient | 1190.48 | 261.39 | 62 | 4.554 | 2.51E-05 |
| Red x Low Nutrient | 365.08 | 261.39 | 62 | 1.397 | 0.167486 |

Table S17: ANOVA summary statistics for the effects of light color, nutrient level, and their interaction on dinoflagellate biomass

| | Sum of Sqs | Mean Sq | Num Df | Den Df | *F* | *p* |
|---|---|---|---|---|---|---|
| Light | 3662132 | 1220711 | 3 | 62 | 7.9407 | 0.000146 |
| Nutrient | 1552154 | 1552154 | 1 | 62 | 10.0967 | 0.002317 |
| Light x Nutrient | 3865079 | 1288360 | 3 | 62 | 8.3807 | 9.25E-05 |

Table S18: ANOVA-like table for the random effects of microcosm replicate and count replicate in dinoflagellate biomass linear mixed effects model

| Model | Number of parameters | log likelihood | AIC | Likelihood ratio | Df | *p* |
|---|---|---|---|---|---|---|
| No random effects | 11 | -482.86 | 987.72 | | | |
| Random effect: Count rep nested in microcosm | 10 | -482.86 | 985.72 | 0 | 1 | 1 |
| Random effect: Microcosm | 10 | -483.67 | 987.35 | 1.629 | 1 | 0.2018 |

Table S19: Output from the green algae biomass linear mixed effects model

| Term | Estimate | Std. Er | df | t | p |
|---|---|---|---|---|---|
| Intercept | 33555.6 | 8321 | 44.1 | 4.033 | 0.000215 |
| Broad | 1851.8 | 11610.9 | 62 | 0.159 | 0.8738 |
| Green | 2821.4 | 11610.9 | 62 | 0.243 | 0.808809 |
| Red | 6063.5 | 11610.9 | 62 | 0.522 | 0.603376 |
| Low Nutrient | -15912.7 | 11610.9 | 62 | -1.37 | 0.175474 |
| Broad x Low Nutrient | -15113.8 | 16420.3 | 62 | -0.92 | 0.360916 |
| Green x Low Nutrient | 8646.8 | 16420.3 | 62 | 0.527 | 0.600355 |
| Red x Low Nutrient | -19182.5 | 16420.3 | 62 | -1.168 | 0.247191 |

Table S20: ANOVA summary statistics for the effects of light color, nutrient level, and their interaction on green algae biomass

| | Sum of Sqs | Mean Sq | Num Df | Den Df | *F* | *p* |
|---|---|---|---|---|---|---|
| Light | 1709125242 | 569708414 | 3 | 62 | 0.9391 | 0.4272798 |
| Nutrient | 8971354405 | 8971354405 | 1 | 62 | 14.7881 | 0.0002863 |
| Light x Nutrient | 2280101479 | 760033826 | 3 | 62 | 1.2528 | 0.298397 |

Table S21: ANOVA-like table for the random effects of microcosm replicate and count replicate in green algae biomass linear mixed effects model

| Model | Number of parameters | log likelihood | AIC | Likelihood ratio | Df | *p* |
|---|---|---|---|---|---|---|
| No random effects | 11 | -746.95 | 1515.9 | | | |
| Random effect: Count rep nested in microcosm | 10 | -746.95 | 1513.9 | 0 | 1 | 1 |
| Random effect: Microcosm | 10 | -746.97 | 1513.9 | 0.039895 | 1 | 0.8417 |

Table S22: Output from the unknown class biomass linear mixed effects model

| Term | Estimate | Std. Er | df | t | p |
|---|---|---|---|---|---|
| Intercept | 3587.3 | 1670.27 | 24.99 | 2.148 | 0.04162 |
| Broad | -3285.71 | 2188.86 | 62 | -1.501 | 0.1384 |
| Green | -3341.27 | 2188.86 | 62 | -1.526 | 0.13197 |
| Red | 7484.13 | 2188.86 | 62 | 3.419 | 0.00112 |
| Low Nutrient | 2222.22 | 2188.86 | 62 | 1.015 | 0.31394 |
| Broad x Low Nutrient | -2507.94 | 3095.51 | 62 | -0.81 | 0.42093 |
| Green x Low Nutrient | -1928.57 | 3095.51 | 62 | -0.623 | 0.53556 |
| Red x Low Nutrient | -3531.75 | 3095.51 | 62 | -1.141 | 0.25829 |

Table S23: ANOVA summary statistics for the effects of light color, nutrient level, and their interaction on unknown class biomass

| | Sum of Sqs | Mean Sq | Num Df | Den Df | *F* | *p* |
|---|---|---|---|---|---|---|
| Light | 1249208050 | 416402683 | 3 | 62 | 19.3137 | 5.87E-09 |
| Nutrient | 953515 | 953515 | 1 | 62 | 0.0442 | 0.8341 |
| Light x Nutrient | 29740930 | 9913643 | 3 | 62 | 0.4598 | 0.7113 |

Table S24: ANOVA-like table for the random effects of microcosm replicate and count replicate in the unknown class biomass linear mixed effects model

| Model | Number of parameters | log likelihood | AIC | Likelihood ratio | Df | *p* |
|---|---|---|---|---|---|---|
| No random effects | 11 | -640.8 | 1303.6 | | | |
| Random effect: Count rep nested in microcosm | 10 | -640.8 | 1301.6 | 0 | 1 | 1 |
| Random effect: Microcosm | 10 | -641.25 | 1302.5 | 0.90031 | 1 | 0.3427 |

Table S25 ANOVA-like table for the random effects of microcosm replicate and count replicate in our filamentous cyanobacteria type linear mixed-effects model.

| Model | Number of parameters | log likelihood | AIC | Likelihood ratio | Df | *p* |
|---|---|---|---|---|---|---|
| No random effects | 11 | -3393.6 | 6809.1 | | | |
| Random effect: Count rep nested in microcosm | 10 | -3393.6 | 6807.1 | 0 | 1 | 1 |
| Random effect: Microcosm | 10 | -3395.4 | 6810.7 | 3.5776 | 1 | 0.05856 |

Table S26: Output from the filamentous cyanobacteria biomass linear mixed effects model

| Term | Estimate | Std. Er | df | t | p |
|---|---|---|---|---|---|
| Intercept | 75.4 | 14331.32 | 13.11 | 0.005 | 0.995882 |
| Broad | 64439.15 | 16990.51 | 266.04 | 3.793 | **0.000185** |
| Green | 5244.05 | 16990.51 | 266.04 | 0.309 | 0.757833 |
| Red | 10396.83 | 16990.51 | 266.04 | 0.612 | 0.541113 |
| Low Nutrient | 214.29 | 16990.51 | 266.04 | 0.013 | 0.989947 |
| Broad x Low Nutrient | -61882.71 | 25636.51 | 267.55 | -2.414 | **0.016457** |
| Green x Low Nutrient | -4105.16 | 24028.2 | 266.04 | -0.171 | 0.864474 |
| Red x Low Nutrient | 5373.02 | 24028.2 | 266.04 | 0.224 | 0.82323 |

Table S27: Output from the non-filamentous cyanobacteria biomass linear mixed effects model

| Term | Estimate | Std. Er | df | t | p |
|---|---|---|---|---|---|
| Intercept | 148.8 | 4604.4 | 35.1 | 0.032 | 0.9744 |
| Broad | 484.8 | 6106.4 | 542.1 | 0.079 | 0.93675 |
| Green | 17043.7 | 6106.4 | 542.1 | 2.791 | **0.00544** |
| Red | 3007.9 | 6106.4 | 542.1 | 0.493 | 0.6225 |
| Low Nutrient | 668.6 | 6106.4 | 542.1 | 0.11 | 0.91285 |
| Broad x Low Nutrient | 11503.4 | 9195.1 | 541.6 | 1.251 | 0.21146 |
| Green x Low Nutrient | 13523.8 | 8635.7 | 542.1 | -1.566 | 0.11792 |
| Red x Low Nutrient | 5136.9 | 8635.7 | 542.1 | 0.595 | 0.5522 |

Table S28: ANOVA-like table for the random effects of microcosm replicate and count replicate in our non-filamentous cyanobacteria type linear mixed-effects model.

| Model | Number of parameters | log likelihood | AIC | Likelihood ratio | Df | *p* |
|---|---|---|---|---|---|---|
| No random effects | 11 | -6506.3 | 13035 | | | |
| Random effect: Count rep nested in microcosm | 10 | -6506.3 | 13033 | 0 | 1 | 1 |
| Random effect: Microcosm | 10 | -6506.7 | 13033 | 0.6752 | 1 | 0.4112 |
